# Constrained optimisation of divisional load in hierarchically-organised tissues during homeostasis

**DOI:** 10.1101/2021.10.07.463365

**Authors:** Peter Ashcroft, Sebastian Bonhoeffer

**Affiliations:** Institute of Integrative Biology, ETH Zurich, Switzerland

**Author notes:** **Code availablity:** Code is publicly available at http://github.com/ashcroftp.

## Abstract

It has been hypothesised that the structure of tissues and the hierarchy of differentiation from stem cell to terminally-differentiated cell play a significant role in reducing the incidence of cancer in that tissue. One specific mechanism by which this risk can be reduced is by minimising the number of divisions – and hence the mutational risk – that cells accumulate as they divide to maintain tissue homeostasis. Here we investigate a mathematical model of cell division in a hierarchical tissue, calculating and minimising the divisional load while constraining parameters such that homeostasis is maintained. We show that the minimal divisional load is achieved by binary division tress with progenitor cells incapable of selfrenewal. Contrary to the protection hypothesis, we find that an increased stem cell turnover can lead to lower divisional load. Furthermore, we find that the optimal tissue structure depends on the time horizon of the duration of homeostasis, with faster stem cell division favoured in short-lived organisms and more progenitor compartments favoured in longer-lived organisms.

## 1 Introduction

Many tissues in the human body undergo constant regeneration of their constituent cells. This regeneration allows the tissue to maintain an effective function and to prevent damage accumulation, for example from exposure to acidic environments in the gut or ultraviolet light on the skin. In organs such as the colon, skin, and hematopoietic system, the regeneration is managed through a hierarchical organisation of stem and progenitor cells. These cells undergo amplification of numbers and phenotypic differentiation to produce the functional, terminally-differentiated cells in large enough quantities to support the functioning of the tissue. The hematopoietic system is the archetypal example of such a tissue: in humans ∼ 10^5^ hematopoietic stem cells (Lee-Six et al., 2018) are ultimately responsible for producing ∼ 10^11^ terminally-differentiated cells per day to satisfy respiratory, immune, and coagulation demands (Kaushansky et al., 2016; Vaziri et al., 1994; Nombela-Arrieta & Manz, 2017).

It has been hypothesised that hierarchical population structures have evolved to protect against the accumulation of mutant cells (Cairns, 1975; Nowak et al., 2003; Pepper et al., 2007; Werner et al., 2013; Rodriguez-Brenes et al., 2013; Derényi & Szöllősi, 2017; Grajzel et al., 2020). Empirical observations give weight to these claims: despite blood cells being over ten times more frequent than any other human cell type (Sender et al., 2016), hematopoietic cancers make up only a small fraction of cancer incidence statistics (Cancer Research UK, 2015). It has been suggested that the protection against cancer is conferred by limiting the cumulative number of cell divisions needed to maintain tissue homeostasis (Derényi & Szöllősi, 2017), as well as ensuring that the majority of cells are transient such that continued cell turnover washes out mutants (Cairns, 1975; Nowak et al., 2003).

Mathematical models in which cells are grouped into homogeneous compartments (so-called compartmental models) have been employed to test different aspects of these cancer prevention hypotheses. With such models it is possible to calculate the number of divisions that cells undergo as they differentiate from stem cells to terminally-differentiated cells (Dingli et al., 2007; Marciniak-Czochra et al., 2009; Rodriguez-Brenes et al., 2013; Derényi & Szöllősi, 2017; Böttcher et al., 2018; Alvarado et al., 2018). The mutation risk is inherently linked to this number of divisions; every DNA replication event carries a risk of mistakes, such that more divisions means a higher risk of genetic damage. Once an expression is obtained for the number of cell divisions across the tissue, it is then possible to optimise the tissue architecture in order to minimise the number of divisions and the risk of mutation accumulation (Derényi & Szöllősi, 2017; Alvarado et al., 2018).

Using a computational model, Pepper et al. (2007) showed that increasing the number of compartments in hierarchically-organised tissues limits the somatic evolution (accumulation of multiple beneficial mutations) of those cells. Similarly, Al-varado et al. (2018) focussed optimising tissue structures to limit mutation accumulation. They found that the optimal tissue structure depends on the objective of optimisation: minimising the number of cells with a single mutation in the tissue requires a high degree of self-renewal across the compartments, while the time until two-hit mutants are generated is maximised by the non-self-renewing binary division tree. Derényi & Szöllősi (2017) were the first to compute the divisional load that is accumulated when producing a lifetime supply of terminally-differentiated cells, rather than explicitly focussing on somatic evolution. They found that optimal tissues which minimise divisional load follow a binary tree structure with no propensity for self-renewal among the progenitor cells, as well as a power law increase in differentiation output along the hierarchy.

What is missing is the following: what happens if we constrain the optimisation of divisional load by ensuring tissue homeostasis, and what changes if we minimise divisions accumulated after stem cell differentiation or if we consider the total divisions (including stem cells) accumulated across a lifetime. Furthermore, what is the impact of the difference between the timescale on which evolution acts (i.e. until reproductive senescence) and the actual lifetime of an individual on divisional load?

In the next section, we introduce our model of cell proliferation in a hierarchical population structure. For the purpose of our analyses, we primarily consider a reduced model in which only symmetric division events can occur. The full model, which considers symmetric and asymmetric cell divisions, as well as cell death, is outlined but the full analysis can only be found in the Appendices. We further consider the possibility for stem cells to be non-dividing, as it has been suggested that quiescence is another mechanism of cancer defence in hierarchical tissues (Adams et al., 2015). Using these models we quantify the number of divisions that cells accumulate. We then use the method of Lagrange multipliers to compute the optimum tissue architecture which minimises divisional load, while constraining the number of terminally-differentiated cells under homeostatic conditions. This optimisation is performed on post-stem cell divisions, as well as for total divisions accumulated over a given time horizon.

## 2 Methods

### 2.1 Model structure

We consider a compartmental model for cell proliferation in a hierarchical tissue, following the likes of Marciniak-Czochra et al. (2009); Rodriguez-Brenes et al. (2011); Stiehl & Marciniak-Czochra (2011); Böttcher et al. (2018); Nienhold et al. (2020). Cells are collected into compartments which represent their level of differentiation. Within each compartment, the cells are indistinguishable. The compartments are labelled by *i* ∈ {1, 2, …, *n*}, where *i* = 1 corresponds to stem cells (SC), 1 < *i* < *n* are progenitor cells, and *i* = *n* are terminally-differentiated cells (TDC). The number of cells in compartment *i* is denoted as *N*_*i*_. Additionally, we consider a compartment of non-dividing (quiescent) SCs, labelled *i* = 0. This structure is highlighted in Figure 1A. We note that we only consider a linear population structure, with population branching here ignored to reduce the number of model parameters.

**Figure 1.**
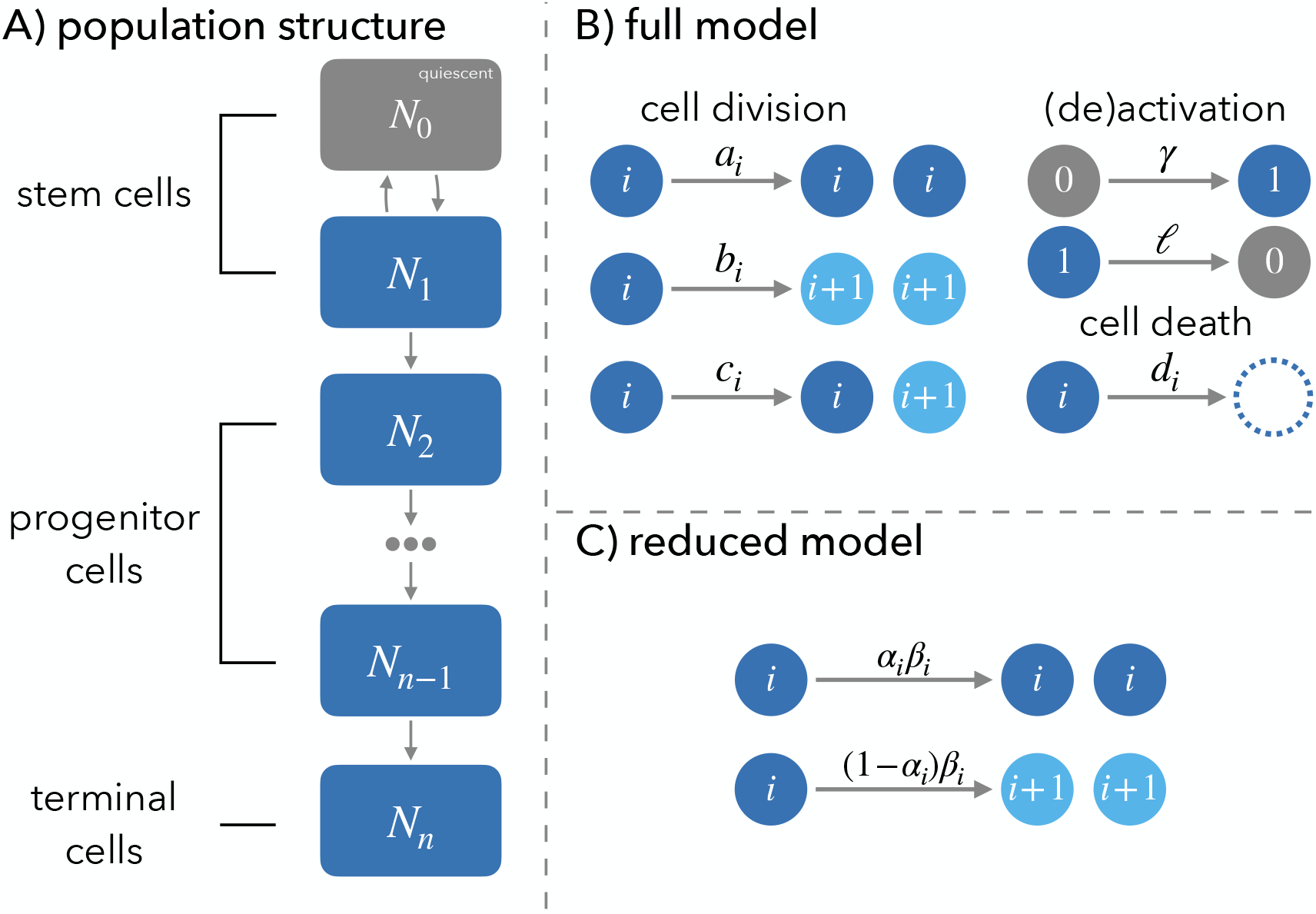
Model schematic. A) We consider a hierarchical population of cells, with stem cells (SC) at the top of the hierarchy giving rise to progenitor cells and ultimately terminally-differentiated cells (TDC). SC can be divided into two subpopulations: a nondividing quiescent fraction (grey) and an active fraction (blue). B) In the full model, cells in compartment *i* ≥1 of the hierarchical population structure can divide symmetrically to generate two offspring of their own type (occurs with rate *a*_*i*_) or two offspring that are differentiated from the parent (rate *b*_*i*_), or they can divide asymmetrically to produce one offspring each type (rate *c*_*i*_). Cells also die at rate *d*_*i*_. Finally, the SC subpopulations are linked through activation and deactivation rates *γ* and *ℓ*, respectively. C) A reduced model is derived from the full model, where cells can only symmetrically self-renew or symmetrically differentiate and there are no quiescent SCs. Here cells in compartment *i* divide with rate *β*_*i*_. With probability *α*_*i*_ the daughters are both of type *i*, and with complimentary probability 1−*α*_*i*_ the daughters are of differentiated type *i* + 1. The parameter *β*_*n*_ assumes the role of the death rate of TDCs, where we have *α*_*n*_ = 0. We also ignore SC quiescence in the reduced model.

### 2.2 Dynamics

Cells move between compartments through differentiation, which is strictly coupled to cell division in this model. A division event produces two daughter cells, each of which may remain in the same compartment as the mother cell (*i*), or differentiate to the next compartment along the hierarchy (*i* + 1). We do not consider the possibility of de-differentiation (i.e. cells cannot move from compartment *i* to compartment *i* − 1). De-differentiation would lead to an increase in divisions accumulated during homeostasis compared to a forward-only differentiating tissue.

Concretely, in the full model (Figure 1B) cells in compartment *i* divide to generate two daughter cells in the same compartment (symmetric self-renewal) with rate *a*_*i*_. With rate *b*_*i*_, two differentiated daughter cells of type *i* + 1 are produced (symmetric differentiation). Cells can also divide asymmetrically with rate *c*_*i*_, such that one daughter remains as type *i* and the other differentiates to type *i* + 1. The quiescent SCs (*i* = 0) do not divide, but can be activated with rate *γ*, while the active SCs (*i* = 1) can deactivate with rate *ℓ*. These dynamics are similar to those proposed by Glauche et al. (2009); Takizawa et al. (2011). Finally, all non-quiescent cells (*i* > 0) die with rate *d*_*i*_.

For the full model (Figure 1B), the number of cells per compartment are described by the following ordinary differential equations (ODE):

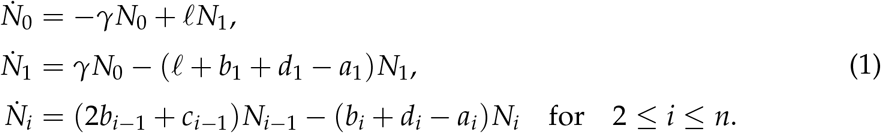

To ensure that the SCs are omnipresent we require *a*_1_ = *b*_1_ + *d*_1_, i.e. any SC lost to differentiation or death is replaced by the self-renewal of another SC. To ensure homoeostasis is possible we require *b*_*i*_ + *d*_*i*_ − *a*_*i*_ > 0 for 2 ≤ *i* ≤ *n*, otherwise *N*_*i*_ would grow exponentially. We further assume that *a*_*n*_ = *b*_*n*_ = *c*_*n*_ = 0 such that TDCs do not divide, as well as *d*_*n*_ > 0 to ensure that a steady-state exists.

Under homeostatic conditions we have 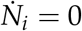 for each compartment 0 ≤ *i* ≤ *n*. From Eq. (1), the equilibrium solution 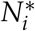 satisfies

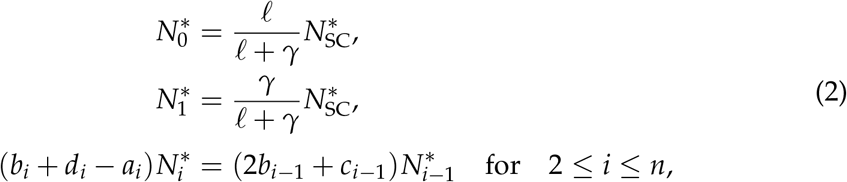

where 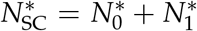 is the total number of stem cells. For 2 ≤ *i* ≤ *n*, Eq. (2) can be solved recursively to give

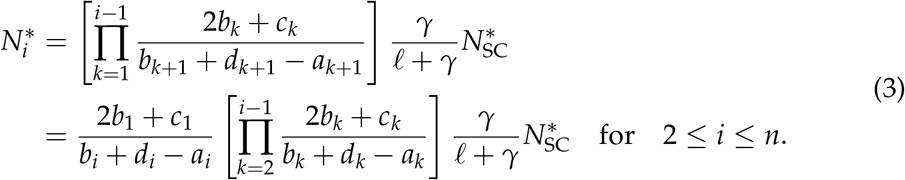

In particular, for TDCs (*i* = *n*) we have

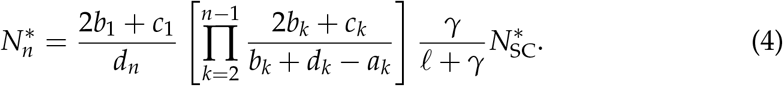

### 2.3 Reduced model

To build intuition and guide analytical calculations, we primarily consider a reduced model (Figure 1C), where we ignore cell death and asymmetric cell division, as well as SC quiescence. In this scenario, we write *a*_*i*_ = *α*_*i*_*β*_*i*_ and *b*_*i*_ = (1 − *α*_*i*_)*β*_*i*_, where *β*_*i*_ is the division rate of cells in compartment *i*, and *α*_*i*_ is their probability of self-renewal (correspondingly, 1 − *α*_*i*_ is the probability of differentiation) following a division event. This reduced model is described by the set of ordinary differential equations

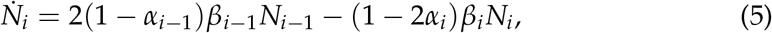

where the first term describes the inflow of two daughter cells from compartment *i* − 1 and the second term is the net loss rate (differentiation minus self-renewal) of cells from compartment *i*. Furthermore, we set *α*_*n*_ = 0 and the parameter *β*_*n*_ assumes the role of the death rate of the TDCs. In the reduced model we ignore SC quiescence by setting *N*_0_ = 0, as well as *γ* = *ℓ* = 0.

Under homeostasis the number of cells per compartment does not change 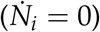. The steady state of Eq. (5), which we write as 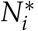, satisfies the relation

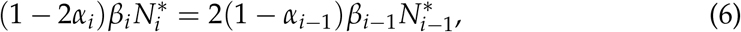

i.e. the net loss from compartment *i* is balanced by the influx from differentiating cells in compartment *i* − 1. For this steady state to exist, we require *α*_1_ = 0.5 and *α*_*i*_ < 0.5 for 1 < *i* < *n*. Solving Eq. (6) recursively, we have

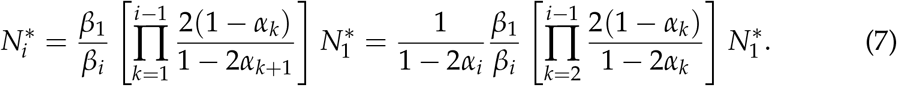

In particular, for TDCs (*i* = *n*) we have

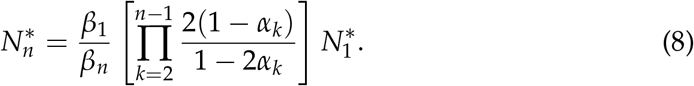

### 2.4 Counting divisions

For compartmental models, the expected number of divisions that cells accumulate in a tissue has previously been calculated using two methods: Firstly, Derényi & Szöllősi (2017) constructed ordinary differential equations (ODE) for the cumulative number of divisions across all cells in a compartment, and from this they extracted the average number of divisions that cells undergo during their lifetime. Alternatively, Böttcher et al. (2018) constructed an explicit set of ODEs for sub-populations of cells of type *i* that have undergone *j* division events, from which the mean number of divisions per cell is easily extracted. These models yield identical results for the mean number of divisions in the full and reduced models, as described in Appendix I and Appendix II, respectively.

Briefly, following the method of Derényi & Szöllősi (2017), the cumulative number of divisions accumulated across all SCs, *T*_1_, in the reduced model satisfies the ODE

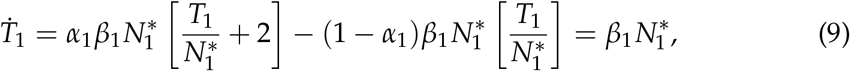

where we have used *α*_1_ = 0.5. Together with the initial condition *T*_1_(0) = 0, Eq. (9) can be solved to give 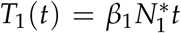. Hence, the average number of divisions per SC at time *t, D*_1_(*t*), satisfies

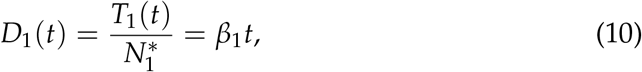

i.e. SC divisions are accumulated linearly with time.

The derivation of the equations for 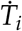 [similar to Eq. (9)] for all compartments 1 ≤ *i* ≤ *n* is described in Appendix I. Ultimately, the expected number of divisions that cells accumulate after a time period *t, D*_*i*_(*t*), is described by a set of linear ODEs which can be solved analytically. However, the resulting expression becomes increasingly complicated for large *n*. Instead, we here make the assumption that the SC dynamics are much slower than non-SCs. We can then express the mean number of divisions per TDC at time *t, D*_*n*_(*t*), as the sum of the SC divisions up to that point in time [Eq. (10)] plus the expected number of divisions that are accumulated *after* SC differentiation, *D*_*n*_. The latter term can be calculated (see Appendix I) as

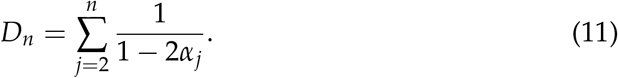

Hence, in the reduced model, the divisions accumulated by TDCs after time *t* is approximated by

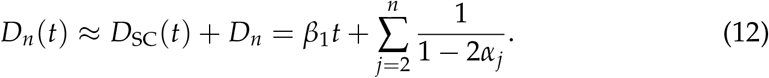

The above methodology can also be applied to the full model, as described in Appendix II. The average number of divisions that a SC (averaged over quiescent and non-quiescent) has accumulated in a time period *t, D*_SC_(*t*), is now

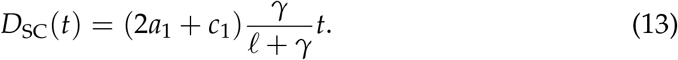

The mean number of divisions that are accumulated by a single TDC after leaving the stem cell compartment, *D*_*n*_, is

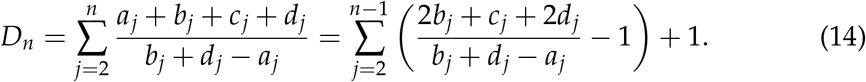

Finally, we approximate the mean number of divisions per TDC at time *t, D*_*n*_(*t*), as the sum of Eq. (13) and Eq. (14):

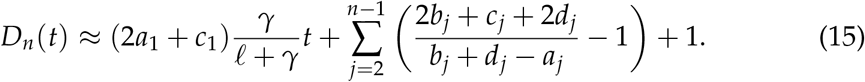

This equation is equivalent to the result of Derényi & Szöllősi (2017), but here with the inclusion of SC quiescence. The calculation differs from Derényi & Szöllősi (2017) in the optimisation procedure in the next steps.

### 2.5 Optimising tissue architecture

Given the expressions for the expected number of divisions accumulated per TDC, we can minimise these values – subject to physical constraints – to deduce the optimal tissue structure parameters which minimise the divisional load. The constraint that we apply is that during homoeostasis the tissue should maintain a given number of TDCs, such that the tissue remains functional. For the reduced model, we derive the constraint from the equilibrium TDC population size, Eq. (8), which we rearrange as

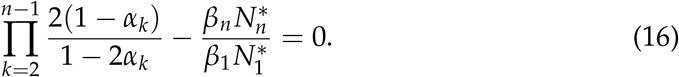

The parameters *α*_*i*_ (1 < *i* < *n*), *β*_1_, 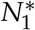 and *n* must satisfy this relation to ensure that enough TDCs are present in the tissue. We keep the number of TDCs 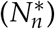 and the death rate of TDCs (*β*_*n*_) fixed, as well as setting *α*_1_ = 0.5 and *α*_*n*_ = 0 as required in the model definition.

We then use the method of Lagrange multipliers (Bertsekas, 2014) to constrain the homeostatic number of TDCs and minimise the tissue’s divisional load. The objective function that is to be optimised is the number of divisions accumulated per TDC, which we can construct in one of two ways: Firstly, we can consider only the divisions accumulated after a SC has differentiated, *D*_*n*_ [Eq. (11)]. In this scenario, we drop the time dependence and only focus on optimising the differentiation structure of the tissue. The Lagrangian function is then constructed as the linear combination of Eq. (11) and Eq. (16), i.e.

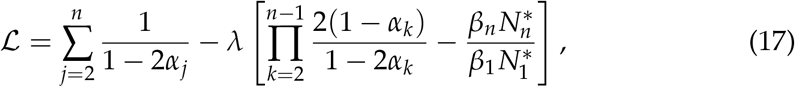

where the coefficient *λ* is the Lagrange multiplier.

Secondly, we can include SC divisional history and ask how many divisions have TDCs accumulated after a given time *t, D*_*n*_(*t*), using the approximation in Eq. (12) as the objective function. The Lagrangian function in this case is

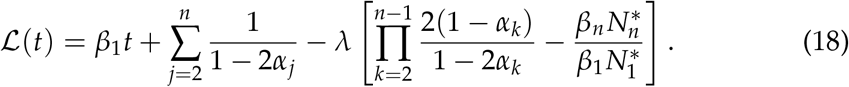

The optimum tissue architecture is then found by computing the stationary points of ℒ or ℒ (*t*) considered as a function of *α*_*i*_ (1 < *i* < *n*), *β*_1_, and the Lagrange multiplier *λ*. Due to the discrete nature of *n*, we cannot identify its optimum value through stationary point analysis. We can, however, identify it through graphical methods. The number of SCs does not appear in the objective function in the Lagrangian, so this too must be optimised by graphical methods.

The Lagrangian functions for the full model can be constructed analogously from the population size constraint in Eq. (4), as well as the number of divisions accumulated after SC differentiation [Eq. (14)] or divisional load after time *t* [Eq. (15)]. See Appendix IV for details.

## 3 Results

Although our results are algebraic, we illustrate these results using the observed parameters of human erythropoiesis (red blood cell production). Concretely, a healthy human has ∼ 10^13^ red blood cells at any given time (Sender et al., 2016) and these cells have a lifetime of ∼ 120 days (Nombela-Arrieta & Manz, 2017). The most recent estimates of human hematopoietic stem cell (HSC) numbers are ∼ 10^5^ cells which divide approximately once per year (Lee-Six et al., 2018).

### 3.1 Optimised tissue architectures in the reduced model

#### 3.1.1 Additional divisions after stem cell differentiation

To determine the architecture that minimises the divisional load, one needs to find the stationary points of the Lagrangian function in Eq. (17). These stationary points satisfy *∂*ℒ / *∂α*_*i*_ = 0 (1 < *i* < *n*) and *∂*ℒ / *∂λ* = 0. As the SC division rate *β*_1_ features only in the constraint and not in the objective function, this parameter cannot be optimised through stationary point analysis. The values of the *α*_*i*_ at the stationary point of ℒ are the parameters which minimise the number of divisions accumulated by the TDCs after leaving the SC compartment, while still maintaining the homeostatic population of TDCs.

From the *∂* ℒ/*∂α*_*i*_ = 0 (for 1 < *i* < *n*) condition, we arrive at

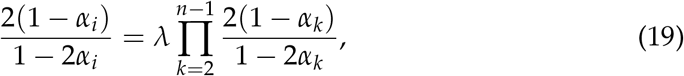

which is only satisfied if

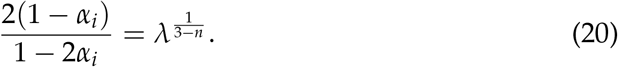

This implies that all *α*_*i*_ are equal for 1 < *i* < *n* in a tissue which minimises the divisional load after SC differentiation. This equivalence of the *α*_*i*_ across compartments was used by Böttcher et al. (2018) and Alvarado et al. (2018) as a simplifying assumption without derivation. Furthermore, this result is consistent with the result of Derényi & Szöllősi (2017), who found that the ratio of differentiation output between two consecutive compartments is constant in optimised tissues.

The value of *λ*, and ultimately the value of the *α*_*i*_, are obtained from the *∂*ℒ /*∂λ* = 0 condition, which is equal to the population size constraint [Eq. (16)]. By substituting Eq. (20) into Eq. (16) and then solving for *λ*, we find the value of the Lagrange multiplier

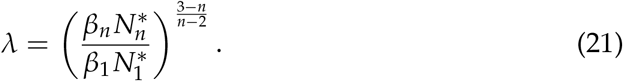

By rearranging Eq. (20) we can now define the values of *α*_*i*_ which minimise the number of divisions per TDC during homeostasis and for a given number of compartments, which we write as 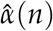:

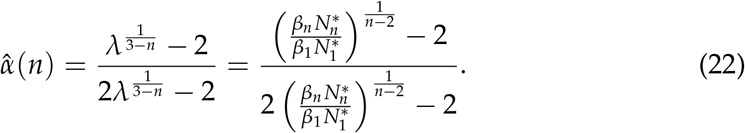

From this equation, we see that an increase in the number of compartments *n* must be compensated by a decrease in the progenitor self-renewal probability 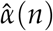 to maintain the homeostatic level of TDCs. However, 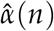 is a probability and is bounded from below by zero. Once *n* is increased sufficiently such that 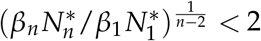, we have to artificially set 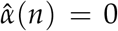 and there will be a new (increased) steady state TDC population size 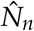. From Eq. (8), the number of TDCs will satisfy

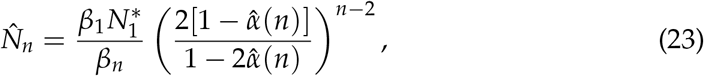

while the number of divisions per TDC after SC differentiation [Eq. (11)] satisfies

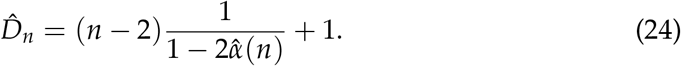

The minimum number of divisions occurs when 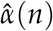 first reaches zero, i.e. when 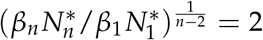. At this point the number of compartments is given by

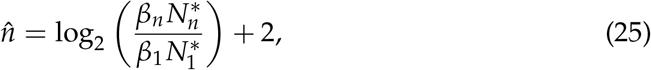

and the divisional load per TDC is

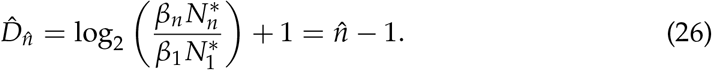

Therefore, perhaps unsurprisingly, the binary division tree 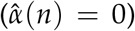 with minimal number of compartments 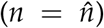 is the tissue structure which minimises divisional load while maintaining the TDC population. As shown by Böttcher et al. (2018), both the mean and variance of the cumulative number of cell divisions is minimal for this non-self-renewing structure. If the number of compartments *n* is smaller than 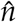, then there must be some self-renewal in the progenitor compartments, which leads to an increase in the cumulative number of divisions. If 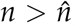, then there are more divisions than required to maintain the TDC population and the tissue produces more TDCs than necessary.

The values of 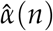 (probability of self-renewal that minimises divisional load per TDC), 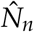 (the corresponding number of TDCs), and 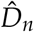 (the divisional load per TDC) are illustrated in Figure 2A-C as a function of total SC turnover 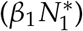 and the number of compartments in the tissue (*n*). The line described by Eq. (25) (dashed diagonal) does indeed follow the minimum divisional load; any change in *either* SC turnover or the number of compartments leads to an increase in divisional load. However, by moving along this line [Eq. (25)], in particular to higher SC turnover and lower number of compartments, the divisional load can be reduced, as shown in Figure 2F. Therefore, the tissues which minimise divisional load after SC differentiation would have very high SC turnover and few differentiation steps.

**Figure 2.**
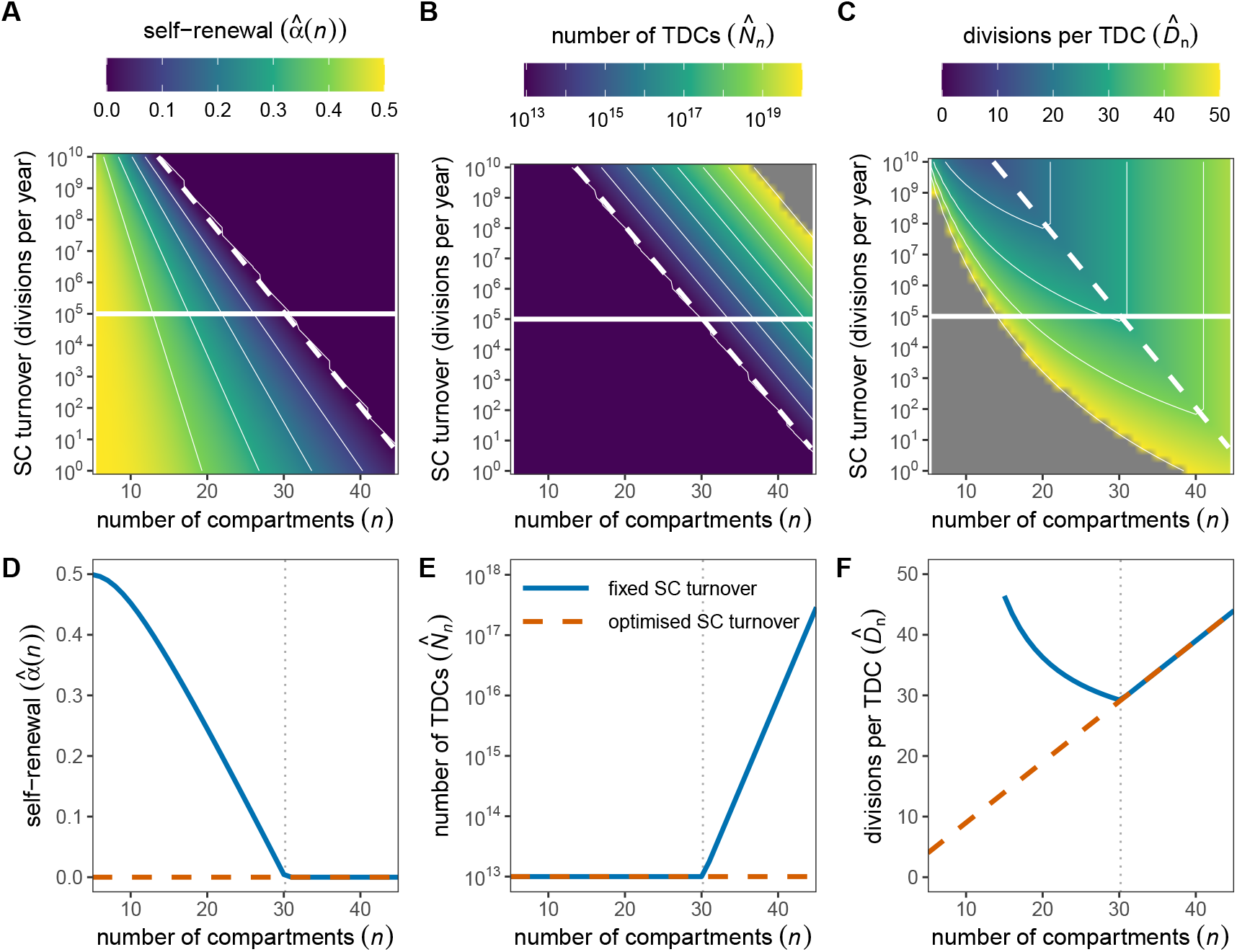
Self-renewal probability (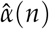; A), TDC population size (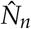; B), and cumulative number of divisions per TDC after SC differentiation (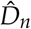; C) as a function of the number of compartments *n* and SC turnover (total divisions per year) 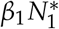 for the reduced model. Here we have fixed the TDC properties *β*_*n*_ = 1/120 day^−1^ and 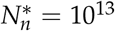, which are approximate numbers for red blood cells in humans (Nombela-Arrieta & Manz, 2017). Horizontal line is the approximate SC turnover in humans (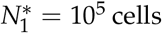 cells; *β*_1_ = 1 yr^−1^; Lee-Six et al. (2018)). Diagonal dashed line represents the optimum number of compartments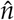 for a given SC turnover which minimises the cumulative number of divisions [Eq. (25)]. The values of 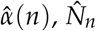, and 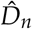 along these slices are shown in panels D, E, and F, respectively. Colour scales for 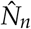 and 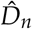 are truncated for visual clarity.

If the SC turnover is fixed for another reason (e.g. due to spatial and nutritional niche competition), then the minimum divisional load occurs when there are enough compartments such that there is no progenitor self-renewal 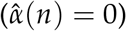, but not too many such that TDCs are not overproduced 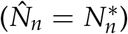 [Figure 2D-F].

#### 3.1.2 Cumulative divisions after lifetime *t*

The objective function in the Lagrangian ℒ(*t*) [Eq. (18)], which now counts SC divisions too, is explicitly dependent on the SC division rate parameter *β*_1_, as well as on the organism’s lifetime. We can now determine the parameters and tissue structure which minimise the divisional load after a time period *τ*, which we call the timescale of optimisation. The optimum parameter values must satisfy *∂*ℒ (*τ*)/*∂α*_*i*_ = 0 (1 < *i* < *n*), *∂*ℒ (*τ*)/*∂λ* = 0, as well as *∂*ℒ (*τ*)/*∂β*_1_ = 0. Note that *τ* is the timescale of optimisation: it is entirely possible that the organism lives for a shorter (*t* < *τ*) or longer (*t* > *τ*) time and we therefore have to consider two timescales.

The partial derivative of ℒ(*τ*) [Eq. (18)] with respect to *α*_*i*_ are unchanged from the previous section, such that the condition Eq. (20) must be satisfied. From the condition *∂*ℒ (*τ*) /*∂β*_1_ = 0, we arrive at

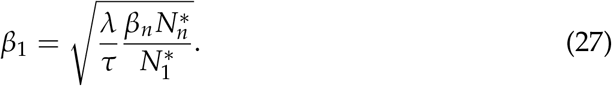

These values of *α*_*i*_ [Eq. (20)] and *β*_1_ [Eq. (27)] can be substituted into the population size constraint Eq. (16), i.e. the *∂*ℒ(*τ*)/*∂λ* = 0 condition, and we can solve for the value of the Lagrange multiplier *λ*:

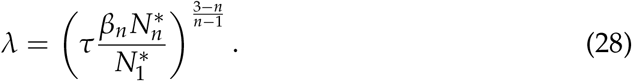

The values of *α*_*i*_ and *β*_1_ which minimise the number of divisions per TDC after time *τ* are thus given by

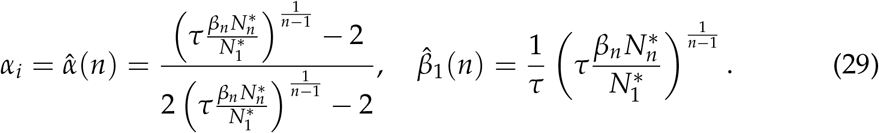

To satisfy 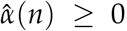, we require 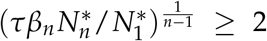. This condition imposes 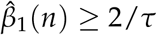.

From Eq. (8), the number of TDCs will satisfy

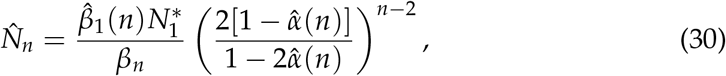

while the number of divisions per TDC after time *t* [Eq. (12)] satisfies

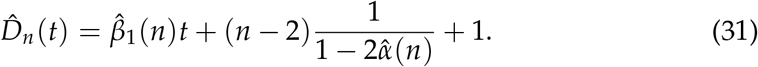

The minimum number of divisions occurs when 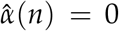and 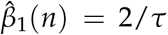, or when 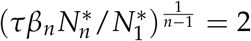. At this point the number of compartments is given by

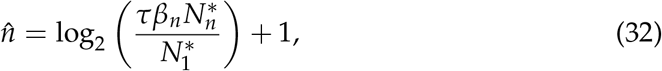

and the divisional load per TDC is

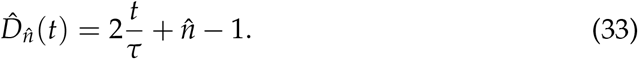

Firstly, we consider the optimised parameters and divisional load when the life-time is equal to the optimisation timescale (*t* = *τ* = 70 years). For this case, the values of 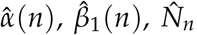, and 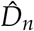 are illustrated in Figure 3A-D as a function of SC number 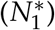 and the number of compartments in the tissue (*n*). The line described by Eq. (32) (dashed diagonal) again follows the minimum divisional load; any change in *either* SC population size or the number of compartments leads to an increase in divisional load. However, by moving along this line [Eq. (32)], in particular to higher SC population size and lower number of compartments, the divisional load per TDC can be reduced, as shown in Figure 3H. Therefore, the tissues which minimise TDC divisional load have many SCs and few differentiation steps. The optimum SC division rate 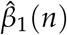 is substantially slower than the observed once per year (Lee-Six et al., 2018), with SCs dividing twice per lifetime in the optimised tissue.

**Figure 3.**
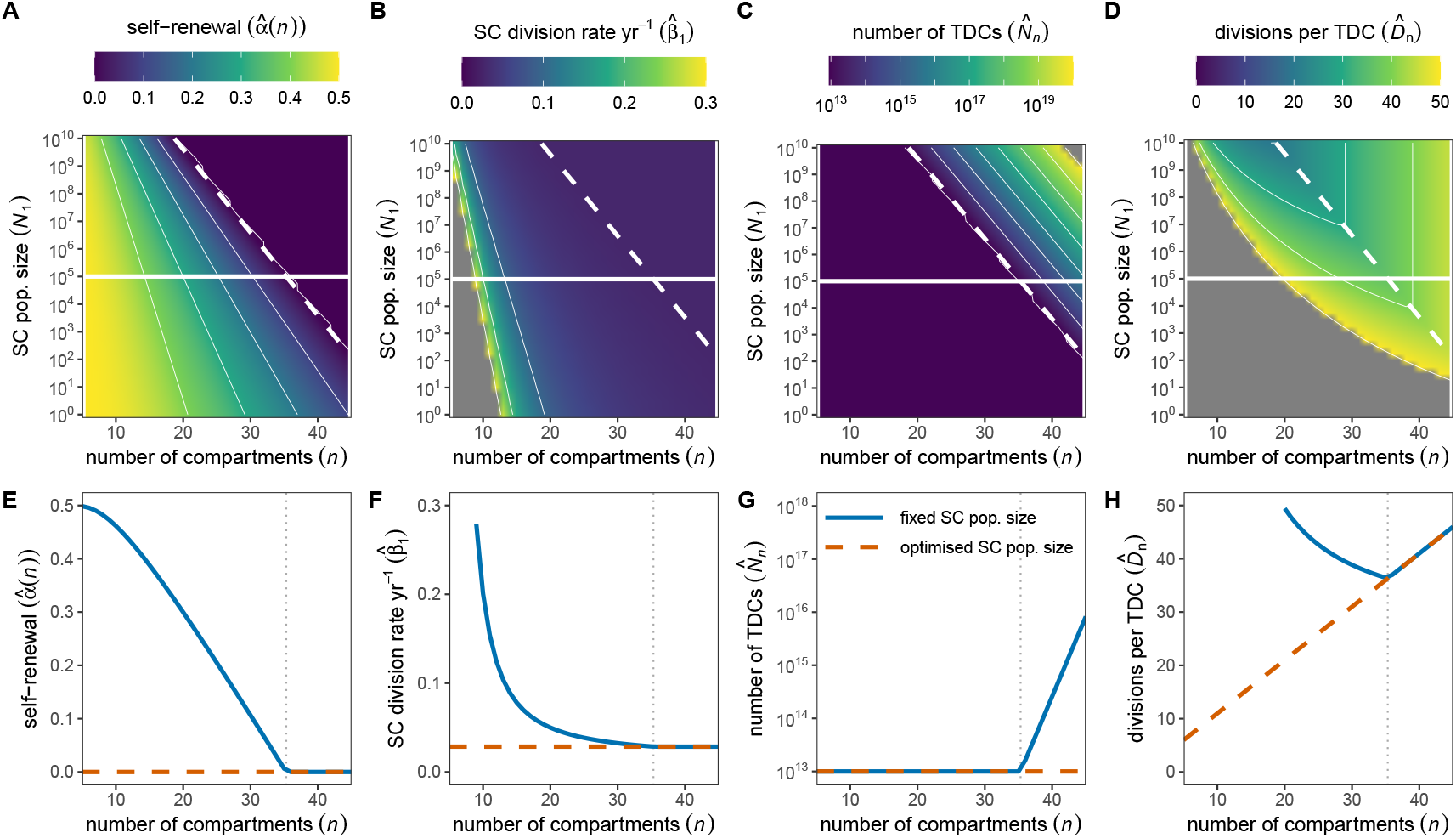
Self-renewal probability (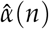: A), SC division rate (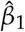; B), TDC population size (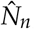; C), and cumulative number of divisions per TDC after time *t* = *τ* = 70 years (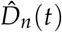; D) as a function of the number of compartments *n* and SC population size 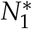 or the reduced model. These parameters/variables are described by Eq. (29), Eq. (30), and Eq. (31). Here we have fixed the TDC properties *β*_*n*_ = 1/120 day^−1^ and 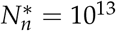 cells, which are approximate numbers for red blood cells in humans (Nombela-Arrieta & Manz, 2017). Horizontal line in A-D is the approximate SC population size in humans (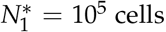; Lee-Six et al. (2018)). Diagonal dashed line represents the optimum number of compartments 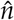 for a given SC population size which minimises the cumulative number of divisions [Eq. (32)]. The values of 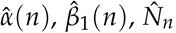, and 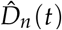 along these slices are shown in panels E-H, respectively. Colour scales for 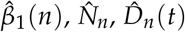 are truncated for visual clarity.

For a fixed SC population size, the SC division rate is a decreasing function of the number of compartments, such that shorter hierarchies require more input from the SC compartment (Figure 3F). The minimum divisional load occurs when there are enough compartments such that there is no progenitor self-renewal 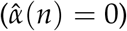, but not too many such that TDCs are not overproduced 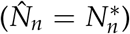, and when SCs are dividing as slowly as possible [Figure 3E-H].

The optimum progenitor self-renewal rate is 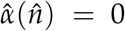. However, the optimum SC division rate 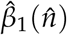, as well as the optimum number of compartments in the hierarchy, 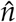, depend on the timescale of optimisation *τ*. As this timescale increases, the optimum SC activity decreases (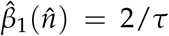; Figure 4A) and the number of amplification steps required to maintain homeostasis increases (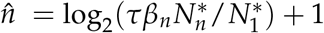; Figure 4B). These properties are independent of the lifetime of the individual. However, the lifetime *t* does contribute to the accumulated divisional load [Eq. (33)]. When we fix *t* = *τ*, we have 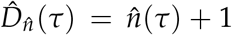, which increases logarithmically with *τ* (blue line in Figure 4C). If the lifetime *t* exceeds the optimisation timescale *τ* (for example when an individual lives beyond reproductive senescence), then TDCs accumulate additional divisions at a rate of 2/*τ* per year (the SC division rate). For short optimisation timescales *τ*, this means a rapid accumulation of additional divisions (orange lines in Figure 4C).

**Figure 4.**
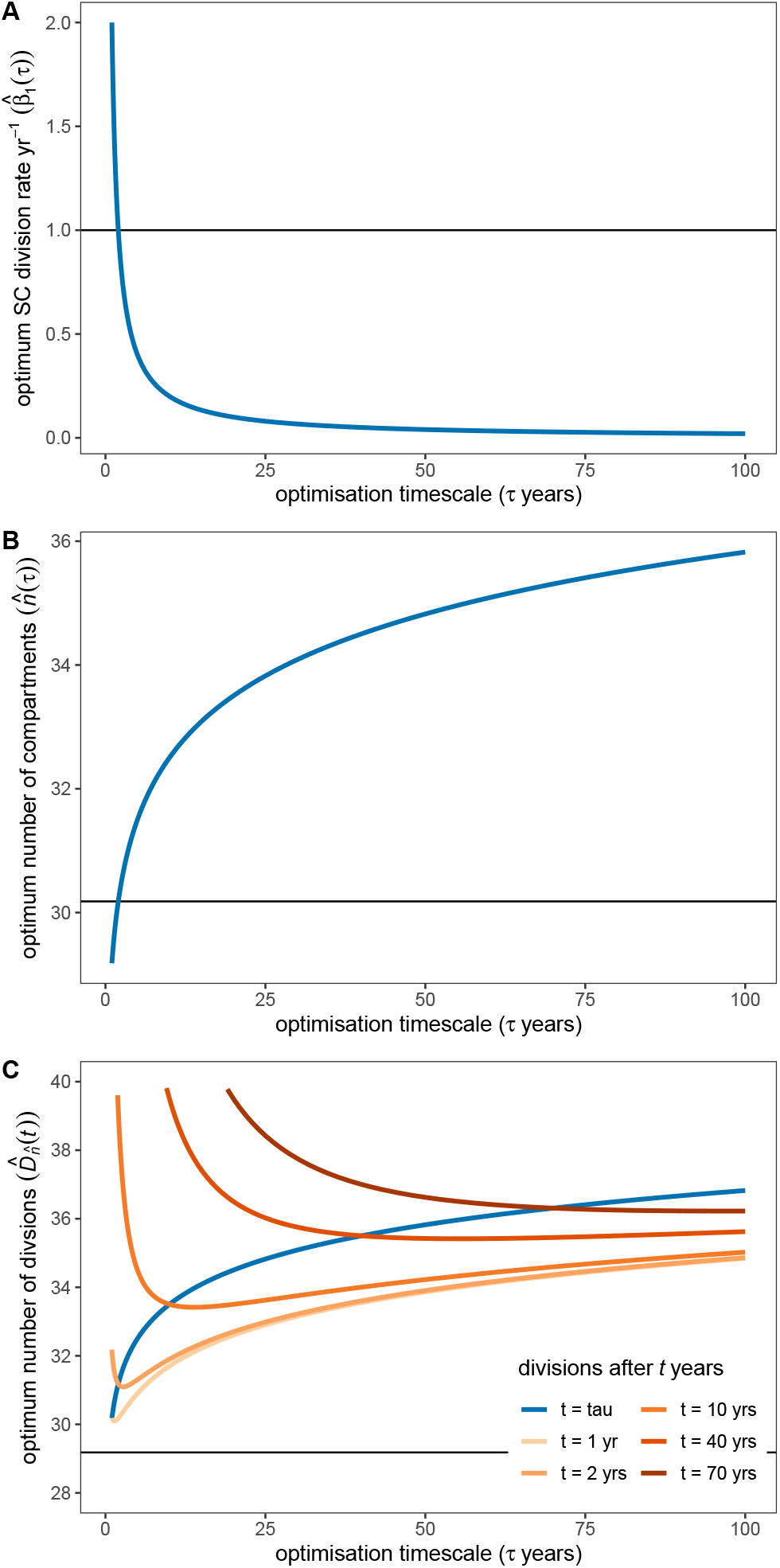
Optimum SC division rate (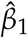; A), number of compartments (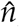; B) and TDC divisional load (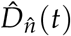; C) as functions of the optimisation timescale *τ*. The optimum self-renewal probability 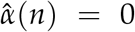 is not shown. Here we have fixed the TDC properties *β*_*n*_ = 1/120 day^−1^ and 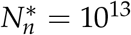 cells, which are approximate numbers for red blood cells in humans (Nombela-Arrieta & Manz, 2017), as well as the SC population size 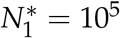 cells (Lee-Six et al., 2018)). Horizontal line in panel A represents the estimated SC division rate in humans (*β*_1_ = 1 yr^−1^; Lee-Six et al. (2018)). Horizontal lines in panels B and C represent the number of compartments and the divisional load when we optimise for divisions accumulated after SC differentiation (i.e. without time).

This rapid accumulation of divisions can be seen more clearly by plotting 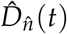 as a function of lifetime *t* for different optimisation timescales *τ* (Figure 5). We also see that choosing an optimisation timescale *τ* that exceeds the lifetime *t* results in more accumulated divisions than necessary in early life, due to the increased number of differentiation steps.

**Figure 5.**
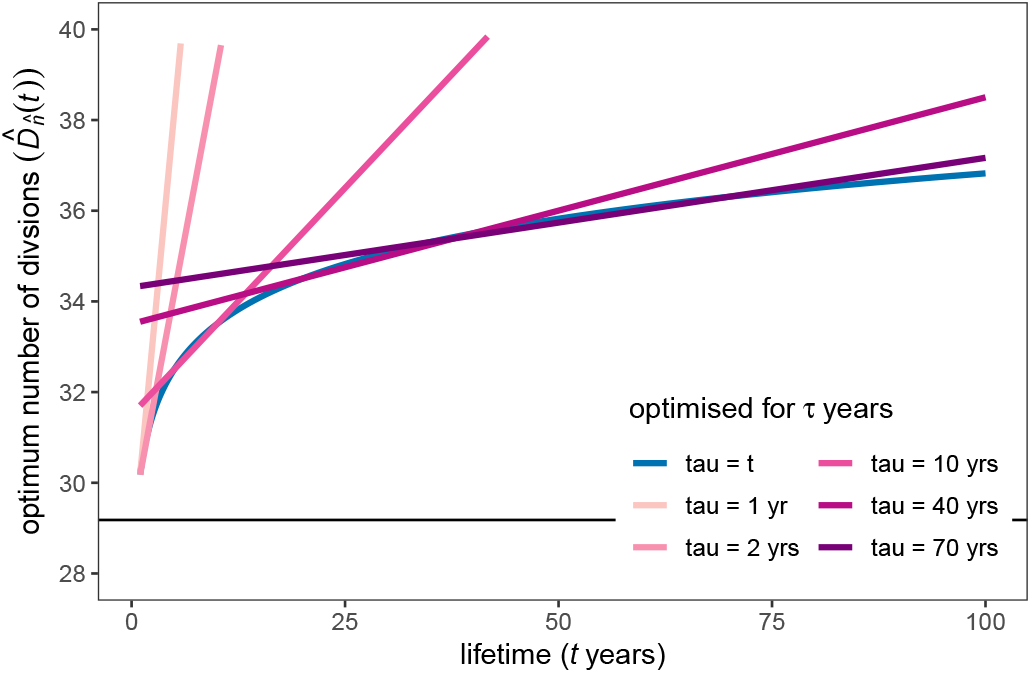
Optimum TDC divisional load 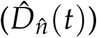 as a function of lifetime *t* for various optimisation timescales *τ* (colours). Here we have fixed the TDC properties *β*_*n*_ = 1/120 day^−1^ and 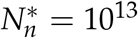 cells, which are approximate numbers for red blood cells in humans (Nombela-Arrieta & Manz, 2017), as well as the SC population size 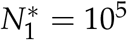 cells (Lee-Six et al., 2018)). Horizontal line representd the TDC divisional load when we optimise for divisions accumulated after SC differentiation (i.e. without time).

### 3.2 Optimised architectures in the full model

For the full model, including quiescence and asymmetric division, we can repeat the above analyses using the Lagrangian function

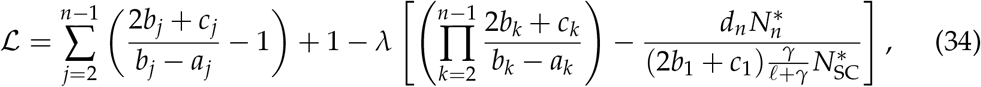

for optimising the divisional load after SC differentiation, and

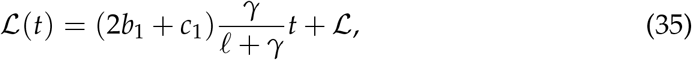

for optimising divisional load after a lifetime *t*. Note that cell death would have to be compensated by additional cell divisions and will result in an increase in the average number of divisions of the surviving cells. We have therefore immediately set *d*_*i*_ = 0 for 1 ≤ *i* < *n*. We have also used the condition *a*_1_ = *b*_1_, which is required for homeostasis.

The optimisation process is described in full in Appendix IV. From this analysis we find that the ratio

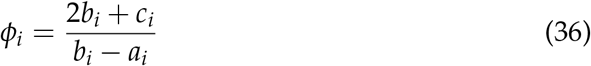

is conserved across all progenitor compartments in optimised tissues, i.e. 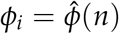 for 1 < *i* < *n*. Because of the conditions *b*_*i*_ − *a*_*i*_ > 0, as well as *a*_*i*_ ≥ 0, *b*_*i*_ ≥ 0, and *c*_*i*_ ≥ 0, this ratio is bounded by 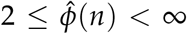. When considering only divisions accumulated after SC differentiation (i.e. without time dependence), this ratio takes the value

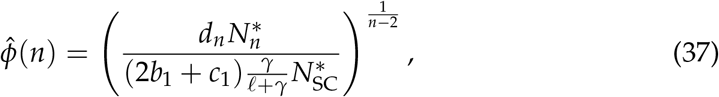

and the divisional load per TDC is

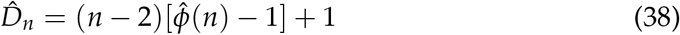

This divisional load is minimised when 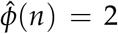, which can only be achieved if *a*_*i*_ = *c*_*i*_ = 0, i.e. progenitor cells (1 < *i* < *n*) do not self-renew or divide asymmetrically, they only differentiate. This pattern was also described by Derényi & Szöllősi (2017). By increasing the SC differentiation flux 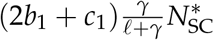, we can reduce the number of compartments while maintaing the TDC population, hence reducing the divisional load (as described for the reduced model).

When we optimise the tissue architecture based on the total TDC divisional load including SC divisions, we have to include an optimisation timescale *τ*. From our analysis (Appendix IV) we find the following relations for the ratio 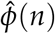 and the total SC division rate:

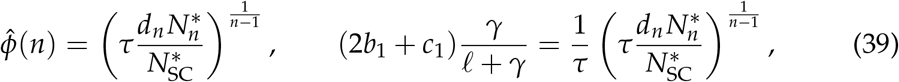

and the divisional load after a lifetime *t* satisfies

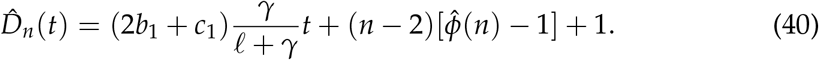

From the relation for the total SC division rate, we observe that there is a trade-off between the division rate of active SCs (2*b*_1_ + *c*_1_) and the fraction of actively dividing SCs [*γ*/(*ℓ* + *γ*)]: If active SCs are dividing faster, then the number of actively dividing cells can be smaller. The total SC division rate is bounded by 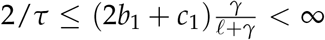. The TDC divisional load is minimised when 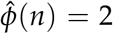, 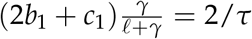 and 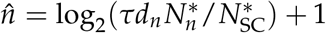.

## 4 Discussion

In this study we set out to quantify and then minimise the divisional load that is accumulated in terminally-differentiated cells (TDC) in a homeostatic tissue. For insight we focussed on a reduced dynamical model in which a division event either lead to symmetric self-renewal or symmetric differentiation. Homeostasis was enforced by constraining the number of progenitor cell compartments (i.e. differentiation steps), self-renewal probabilities per compartment, and division rates per compartment such that the TDC population remains at its steady-state level.

In tissues which minimise the divisional load, we find that the self-renewal probability is constant throughout all progenitor cell compartments. The optimum number of compartments is then the smallest number for which progenitor self-renewal can be zero without producing excess TDCs. Therefore, the optimum tissue structure which minimises divisional load is a binary division tree. By increasing SC turnover, either by increasing SC numbers or SC division rate, the number of progenitor compartments that are required to amplify cell numbers can be reduced and subsequently the divisional load of TDCs is also reduced. Therefore, if optimising purely based on divisional load as we have done here, it would make sense to have a high-turnover SC compartment, which is the opposite of what we actually observe in, e.g., the hematopoietic system or colonic crypts. Hence, there must be other selection pressures or physiological constraints at play that result in the observed low SC turnover in human tissues.

The above results were obtained based on the divisional load accumulated in the tissue after SC differentiation and in the absence of a time component. However, in our model, SCs accumulate divisions linearly with time. When including these SC divisions, we find that in the short term it is better to have higher SC turnover and fewer progenitor compartments, but in the long term having more compartments and less SC turnover is optimal. Therefore, based on our optimisation criteria, we would expect that in longer lived organisms (with the same homeostatic TDC number) SCs should divide more slowly and there should be a greater number of amplification steps to minimise division accumulation. A recent measurement of somatic mutation rate across animals with different lifespans supports this hypothesis, with longer lived animals having a lower rate of somatic mutation accumulation (Cagan et al., 2021). Furthermore, we find that underestimating the timescale of optimisation relative to lifetime leads to a large increase in divisional load at ages beyond the optimisation timescale.

Our full model of cell dynamics also allows asymmetric division, cell death, and SC quiescence. Repeating the above analysis, we find that the ratio of differentiation output to net loss in a progenitor compartment is conserved across progenitor compartments in optimised tissues. This is a more general principle than derived by Derényi & Szöllősi (2017) who found that the ratio of differentiation output between two consecutive progenitor compartments is constant in optimised tissues. Furthermore, the minimum divisional load is achieved when the differentiation output to net loss ratio is equal to two, i.e. when progenitor cells do not self-renew and differentiate only symmetrically. This is something that could be compared to empirical data to check for tissue optimisation, if such data existed. Finally, from our model and analysis we find that there is a trade-off between the SC division rate and the fraction of quiescent SCs: faster dividing SCs would require a larger quiescent fraction to maintain the minimum divisional load (for a given number of progenitor compartments).

Interestingly, although optimal tissues have a constant self-renewal across progenitor cells, the division rates of the progenitor cells do not affect the divisional load and are thus unconstrained by our optimisation procedure. Briefly, the progenitor compartment size 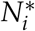 is inversely proportional to the division rate in that compartment [Eq. (7)], so the total differentiation flux out of that compartment 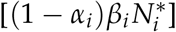 is independent of *β*_*i*_. Therefore, the relationship between compartment sizes in these optimised tissues are unconstrained as well: I.e., it is not necessary to have a monotonically increasing compartment size for the tissue to minimise divisional load. In optimised tissue structures (binary division trees), we do have a doubling of the flux of cells between consecutive progenitor compartments. By assuming a constant division rate across progenitor compartments, we arrive at the doubling of subsequent progenitor compartment sizes in these optimised tissues.

Our conclusions are in agreement with those of Derényi & Szöllősi (2017), despite different optimisation procedures: We both find that hierarchically-organised tissues can be tuned to minimise divisional loads, and therefore limit the occurrence of somatic mutations during homeostasis. Beyond Derényi & Szöllősi (2017), we show that the constant ratio of differentiation outputs between subsequent compartments can be generalised further: we find that the ratio of differentiation output to net loss per compartment is conserved across all compartments, which in the absence of asymmetric divisions means that self-renewal probability is constant across all compartments. Furthermore, we explore the impacts of varying the optimisation timescale, lifetime, SC numbers, and SC quiescence.

Although the number of divisions itself is an important measure for damage accumulation per cell, it is not the full story in terms of cancer prevention. Nowak et al. (2003) formalised the concepts put forward by Cairns (1975) to show that preventing self-renewal in all cells downstream of the SC minimises the rate that mutations fixate in the population. This simple so-called “linear model” of somatic evolution corresponds to our optimised binary division tree. Later, Pepper et al. (2007) extended the serial differentiation model to include cell compartments (as in our model). They showed that non-self-renewing tissue structures can suppress cell level selection and somatic evolution, as serial differentiation makes it possible to segregate proliferative activity and population self-renewal into different cell compartments, such that no compartment possesses all the attributes necessary for somatic evolution. Any progenitor self-renewal, including asymmetric divisions, would allow rapid somatic evolution because a mutant clone can sweep to fixation within a population of such cells. Therefore, the tissue structures which we find limit the divisional load also limit the rate of somatic evolution.

Recently, Alvarado et al. (2018) utilised an alternative “top-down” modelling approach in which cells that are lost from the TDC compartment are replaced by pulling from the previous compartment (rather than SC division pushing cells into the next compartment). They showed that by increasing progenitor self-renewal it becomes unlikely that mutations are acquired in the more primitive compartments, but mutations that do occur take longer to be flushed out of the tissue. Overall, large values of progenitor self-renewal result in a smaller number of single-hit mutants across all compartments. However, from the point of view of two-hit mutant generation (e.g. the inactivation of a tumour suppressor gene), less progenitor selfrenewal is advantageous as the lifetime of the transient cells is shorter, and they therefore have less time to accumulate two mutations. This optimal architecture for delaying two-hit mutations corresponds to the binary division tree which limits divisional load. Interestingly, Alvarado et al. (2018) found that the arrangement of progenitor compartment sizes influences the total number of mutants and the rate at which two-hit mutants are generated. This is seemingly in contrast with our observation that progenitor compartment sizes are essentially a free variable and unconstrained by minimising the divisional load. This discrepancy can be explained by the fact that the cell division rates in Alvarado et al. (2018) are determined by a feedback loop and are intrinsically coupled to the self-renewal probabilities.

In our study we have quantified only the mean divisional load per TDC. Böttcher et al. (2018) showed that, based on telomere length distributions, there is significant heterogeneity among divisional loads. In their analysis, Böttcher et al. (2018) showed that not only the mean, but also the variance, increases with increasing progenitor self-renewal, leading to few cells with very high divisional load. This again suggests that the non-self-renewing tissue would be the optimum for minimising the probability of a cell evolving an oncogenic phenotype through mutation accumulation.

Our analysis relies on assumptions about the system being in steady state and parameter values being constant (for example, constant HSC activation and deactivation throughout life). This, however, is not representative of early development when SCs undergo significant expansion and tissue compartments are populated. An expansion phase would require an initially increased SC self-renewal rate to allow the SCs to reach the homeostatic level and to populate the progenitor and TDC compartments.

With this study we have shown that limiting progenitor self-renewal in hierarchical tissues results in the lowest divisional load during homeostatic tissue maintenance. This optimised tissue also satisfies some other somatic evolution constraints put forward by multiple authors. Going forward, we would like to bring all of these aspects together to achieve an overall understanding of the selection pressures that act on and shape tissue organisation.

## Appendix

### Appendix I Counting divisions in the reduced model

#### Total divisional load

We here recall the argument of Derényi & Szöllősi (2017), which we apply to our reduced hierarchical population model. To this end, let *T*_*i*_ be the cumulative total of cell divisions that all of the cells in compartment *i* have undergone. Each individual division event modifies the *T*_*i*_, so we want to construct an equation for this quantity:

1. Following a self-renewal event in compartment *i*, which occurs at a rate *α*_*i*_*β*_*i*_ per cell, the division total in compartment *i* increases. The mother cell had accumulated, on average, *T*_*i*_/*N*^*^, and then generated two daughter cells with *T*_*i*_/*N*^*^ + 1 divisions each. Therefore, the increase in *T*_*i*_ per self-renewal event is 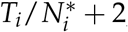;
2. Following a differentiation event in compartment *i* − 1, which occurs at a rate (1 − *α*_*i*−1_)*β*_*i*−1_ per cell, the division total of compartment *i* − 1 decreases (on average) by 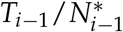. Meanwhile, the division total in compartment *I* increases by 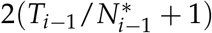, where the factor two reflects the fact that two differentiated daughters are generated per event.

Combining these events, we have the following ODE for the cumulative number of divisions of all cells in compartment *i*:

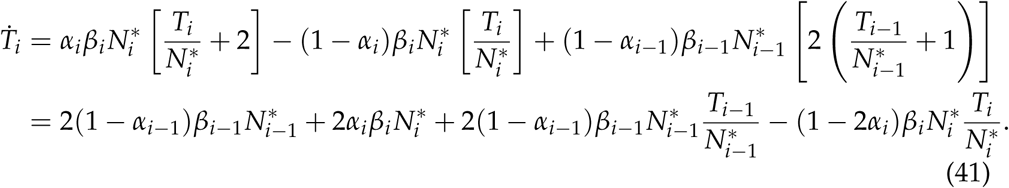

Using the steady-state relation 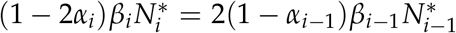 [Eq. (6) in the main text], we recover

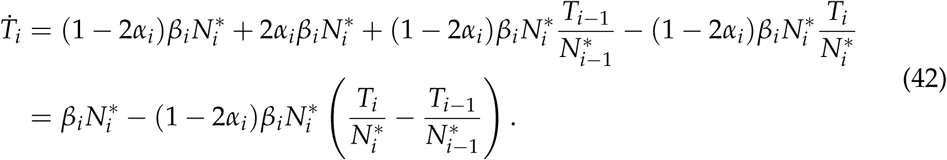

Now writing 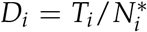 for the average number of divisions per cell in compartment *i*, we have the equation

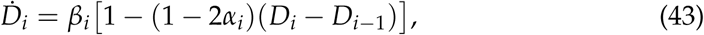

which we can solve recursively. For *i* = 1, we have (using *α*_1_ = 0.5)

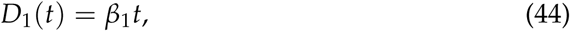

i.e. stem cells accumulate divisions linearly with time.

For the average number of divisions beyond the stem cells, we can look at the steady state of Eq. (43) and set *D*_1_ = 0. From this steady state we have

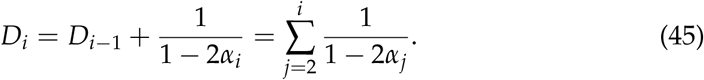

##### Tracking divisions with subpopulations

The number of divisions that a cell goes through can also be computed according to the method of Böttcher et al. (2018). In summary, the number of cells in compartment *i* that have undergone *j* cell divisions follows the ODE

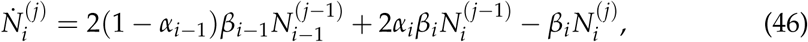

where the three terms correspond to differentiation into compartment *i* from *i* − 1, self-renewal within compartment *i*, and the division of cells which have already accumulated *j* divisions in compartment *i*. To compute the number of divisions a cell undergoes in addition to those in the stem cell compartment (*i* = 1) we set 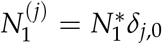, where *δ*_*j*,0_ is the Kroenecker-*δ*, which has value one if *j* = 0 and zero otherwise. We then look at the steady state of Eq. (46), i.e.

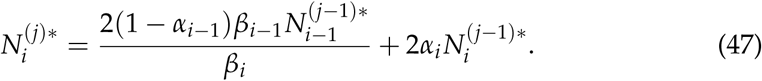

Using the steady-state relationship 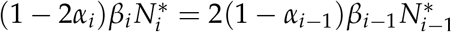 [Eq. (6) in the main text], and defining 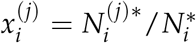(i.e. the fraction of cells in compartment *i* that have undergone *j* divisions), we have

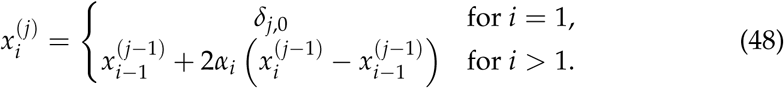

From this, we can compute the mean number of divisions per compartment 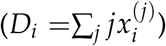 which gives

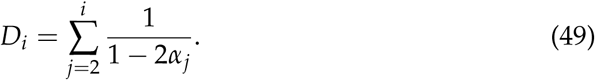

This is identical to Eq. (45).

### Appendix II Counting divisions in the full model

#### Total divisional load

We here again recall the argument of Derényi & Szöllősi (2017), which we apply to our full hierarchical population model. To this end, let *T*_*i*_ be the cumulative total of cell divisions that all of the cells in compartment *i* have undergone. The individual events modify the *T*_*i*_, so we want to construct an equation for this quantity. Firstly, for the cumulative number of divisions of cells in the quiescent compartment *i* = 0 we have

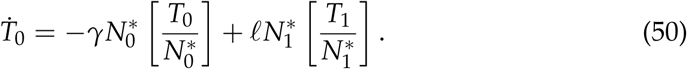

Cells in the active stem cell compartment *i* = 1 can move to and from the quiescent compartment, but can also increase the number of divisions through self-renewal and asymmetric divisions and decrease the number of divisions through differentiation and death. We therefore have

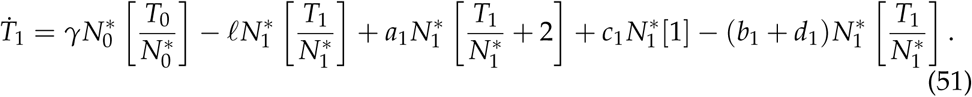

For cells in compartments 2 ≤ *i* < *n* we have the following events which can modify the cumulative number of divisions per compartment:

1. Following a symmetric self-renewal event in compartment *i*, which occurs at a rate *a*_*i*_ per cell, the division total in compartment *i* increases. The mother had accumulated, on average, *T*_*i*_/*N*^*^, and then generated two daughter cells with *T*_*i*_/*N*^*^ + 1 divisions each. Therefore, the increase in *T*_*i*_ per self-renewal event is 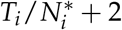.
2. Following a symmetric differentiation event in compartment *i* − 1, which occurs at a rate *b*_*i*−1_ per cell, the division total of compartment *i* − 1 decreases (on average) by 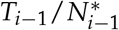. Meanwhile, the division total in compartment *I* increases by 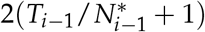, where the factor two reflects the fact that two differentiated daughters are generated per event.
3. Following an asymmetric division event in compartment *i*, which occurs at a rate *c*_*i*_ per cell, the division total in compartment *i* increases by one, and the division total in compartment *i* + 1 increases (on average) by 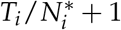.
4. Finally, a death event in compartment *i* occurs at a rate *d*_*i*_ per cell and removes (on average) 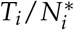 cumulative divisions from that compartment.

Combining these events, we have the following ODE for the cumulative number of divisions of cells in compartment 2 ≤ *i* < *n*:

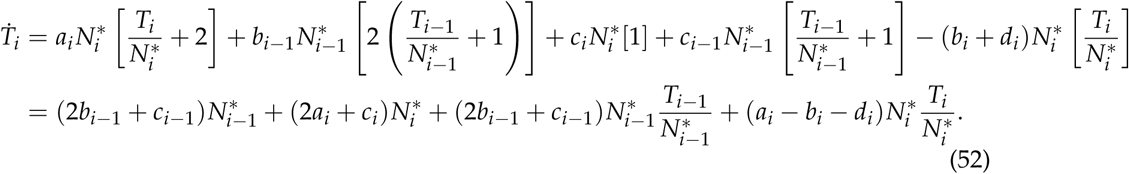

Using the steady-state relation 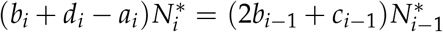 [Eq. (2) in the main text], we recover

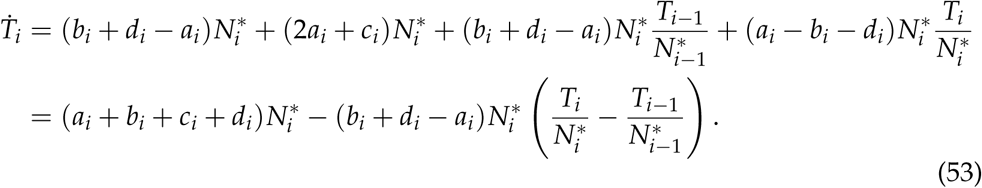

Now writing 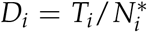 for the average number of divisions per cell in compartment *i*, we have the equations

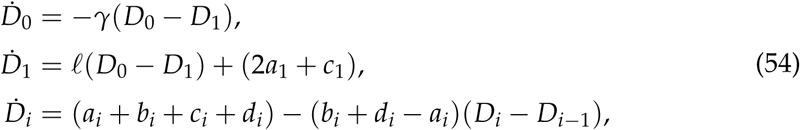

where we have used *a*_1_ = *b*_1_ + *d*_1_ and 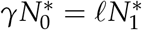. The ODEs for *D*_0_ and *D*_1_ can be solved together [with *D*_0_(0) = *D*_1_(0) = 0] to give

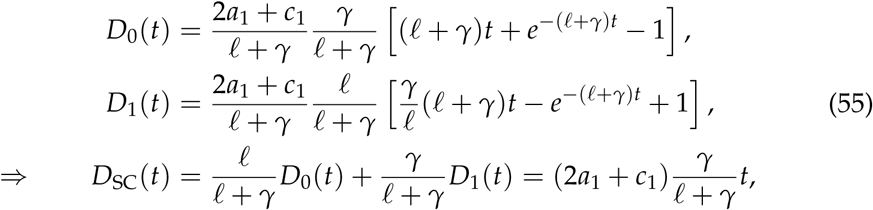

where *D*_SC_(*t*) is the average number of divisions per stem cell across the active or quiescent stem cells. For the average number of divisions beyond the stem cells, we can look at the steady state of Eq. (54) and set *D*_1_ = 0. From this steady state we have

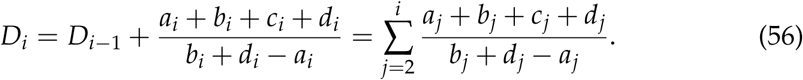

#### Tracking divisions with subpopulations

The number of divisions a cell goes through can also be computed according to the method of Böttcher et al. (2018). In summary, the number of cells in compartment *i* that have undergone *j* cell divisions follow the ODEs

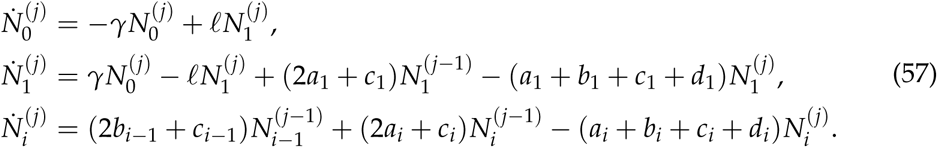

For the stem cells we define 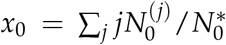 and 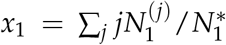 as the mean number of divisions per cell in the quiescent and active compartment. These variables then follow the ODEs

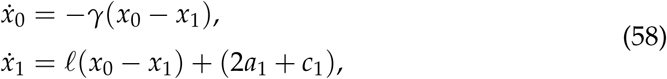

where we have used 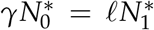 and *a*_1_ = *b*_1_ + *d*_1_. These ODEs can be solved together [with *x*_0_(0) = *x*_1_(0) = 0] to give

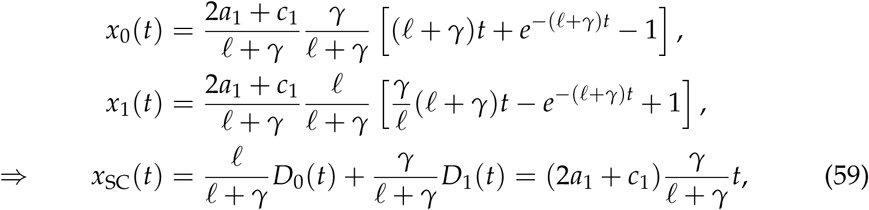

where *x*_SC_(*t*) is the average number of divisions per stem cell ignoring the active or quiescent status.

To compute the number of divisions a cell undergoes in addition to those in the stem cell compartments we set 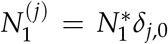. We then look at the steady state of Eq. (57), i.e.

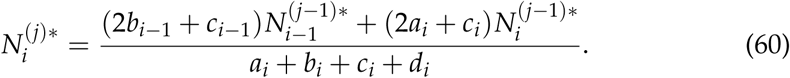

Using the steady-state relation 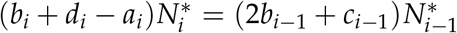 [Eq. (2) in the main text], and defining 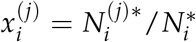 (i.e. the fraction of cells of type *i* that have undergone *j* divisions), we have

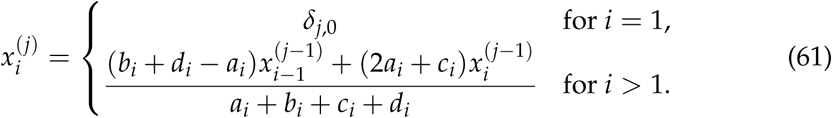

From this, we can compute the mean number of divisions per compartment 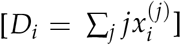 which gives

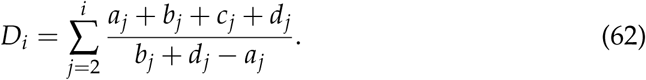

This is identical to Eq. (56).

### Appendix III Tissue optimisation in the reduced model

#### Minimizing additional divisions after stem cell differentiation

Here we optimise the number of divisions that TDCs accumulate after leaving the SC compartment. From Eq. (45), the number of divisions post SC differentiation is

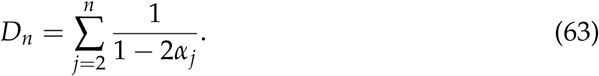

Concretely, we want to solve

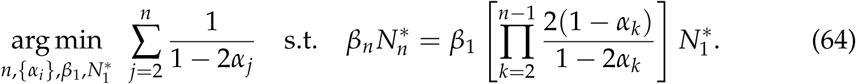

As the parameters *β*_1_ and 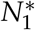 do not appear in the objective function (only in the constraint), they cannot be optimised here. Furthermore, the number of compartments *n* will be optimised through graphical methods due to its discrete nature. Hence, we first identify the parameter set {*α*_*i*_}_1<*i*<*n*_ which minimises the number of divisions.

Following the method of Lagrange, we find the stationary points of the function

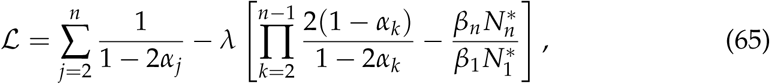

which combines the cumulative number of divisions with the population size constraint. The coefficient *λ* is the Lagrange multiplier, which is to be determined. The stationary points of the Lagrangian function are found by solving *∂*ℒ/*∂α*_*i*_ = 0 and *∂*ℒ/*∂λ* = 0. From the first set of partial derivatives, we arrive at

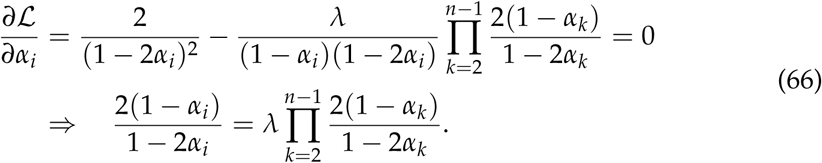

We can also write this as

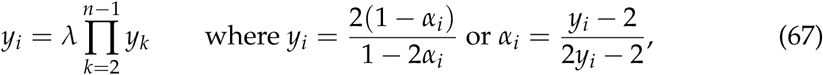

from which we see that all *y*_*i*_ are equal (for 1 < *i* < *n*). Solving this equation, we therefore have

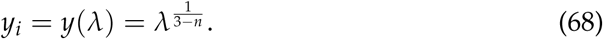

The value of *λ* is obtained from the *∂*ℒ/*∂λ* = 0 equation, i.e. the population size constraint:

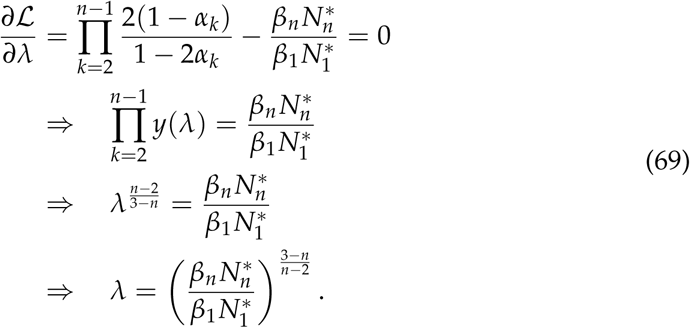

Hence the values of *α*_*i*_ for the *n*-compartment model are given as

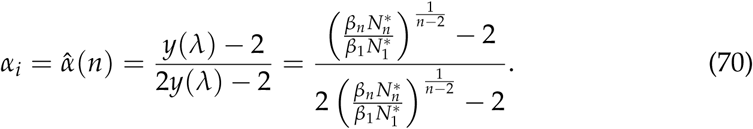

Inserting this value of 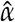 into the expression for the number of divisions per TDC after SC differentiation [Eq. (63)], we arrive at

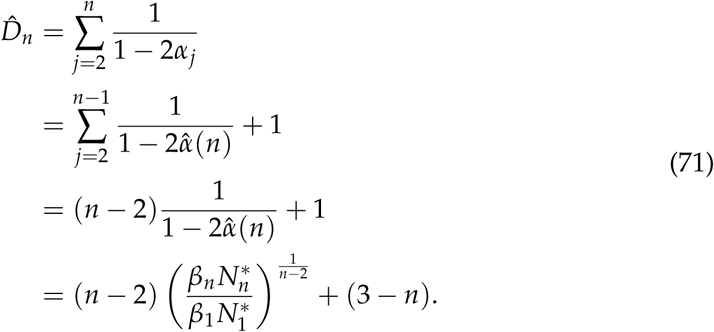

Finally, we note that 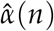 has to bounded below by zero 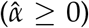. This value is achieved when 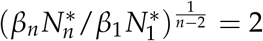 [see Eq. (70)], or when

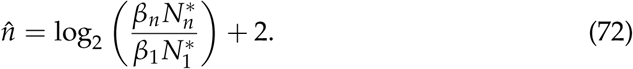

Through graphical methods, we show that 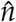 is the optimum number of compartments to minimise the divisional load, where we have 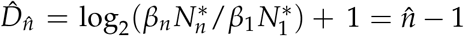.

#### Minimizing cumulative divisions after lifetime *t*

For the number of divisions accumulated after a lifetime *t*, we use the assumption that SC divisions occur much more slowly than non-SCs, such that the number of divisions per TDC at time *t* is approximated by the number of SC divisions at time *t* plus the number of divisions accumulated after SC differentiation. I.e. the number of divisions per TDC at time *t* is

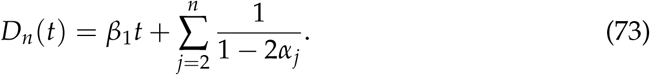

Here we optimise the number of divisions that TDCs will have accumulated after SCs have been dividing for time period *τ*. Note that *τ* is the timescale of optimisation, but the lifetime *t* may be longer or shorter than *τ*. Concretely, we want to solve

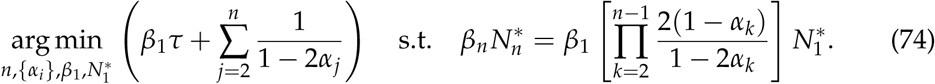

The number of stem cells, 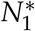, does not appear in the objective function (only in the constraint) and cannot be optimised here. Furthermore, the number of compartments *n* will be optimised through graphical methods due to its discrete nature. Hence, we first identify the parameter set {*α*_*i*_}_1<*i*<*n*_ and *β*_1_ which minimise the divisional load.

Following the method of Lagrange, we find the stationary points of the function

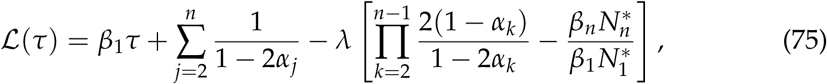

which combines the cumulative number of divisions per TDC with the population size constraint. The stationary points of the Lagrangian function are found by solving ∂ℒ(*τ*)/*∂α*_*i*_ = 0, *∂*ℒ(*τ*)/*∂β*_1_ = 0, and *∂*ℒ(*τ*)/*∂λ* = 0.

The partial derivatives with respect to *α*_*i*_ are equivalent to the previous section, such that we arrive at

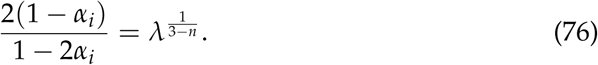

For the partial derivative with respect to *β*_1_, we arrive at

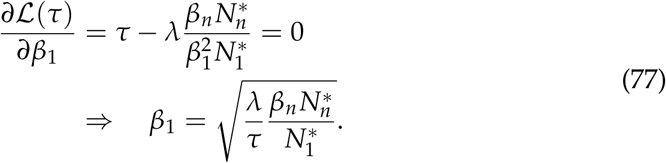

The value of *λ* is obtained from the *∂*ℒ(*τ*)/*∂λ* = 0 equation, i.e. the population size constraint:

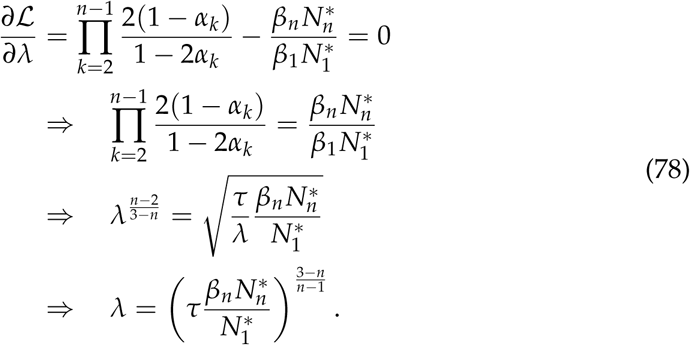

Hence the optimum values of *α*_*i*_ and *β*_1_ are given as

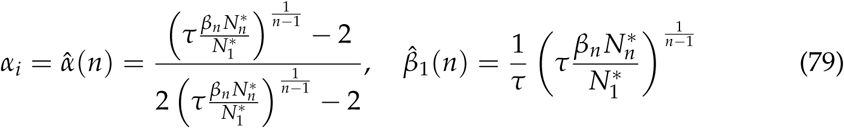

Inserting these values of 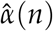 and 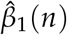 into the expression for the number of divisions per TDC after a lifetime *t* [Eq. (73)], we arrive at

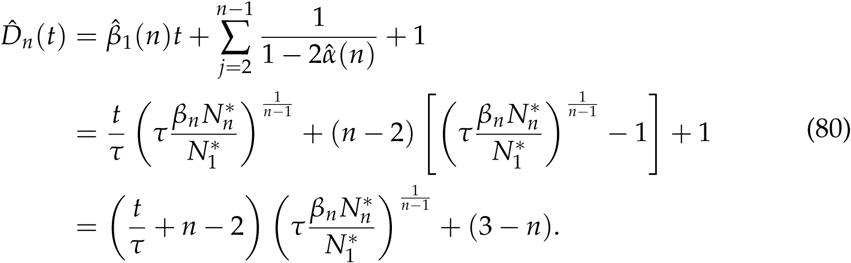

Finally, we note that 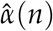 has to bounded below by zero 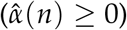. This value is achieved when 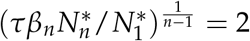[see Eq. (79)], or when

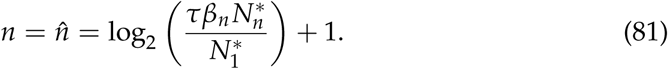

### Appendix IV Tissue optimisation in the full model

#### Minimizing additional divisions after stem cell differentiation

We can repeat the above analysis for the full model, where we want to minimise the number of divisions accumulated by a TDC after leaving the stem cell compartment, i.e. Eq. (56). We first note that cell death would have to be compensated by additional cell divisions and will result in an increase in the average number of divisions of the surviving cells. We can therefore immediately set *d*_*i*_ = 0 for 1 ≤ *i* < *n*. We now want to find the stationary points of the Lagrangian function

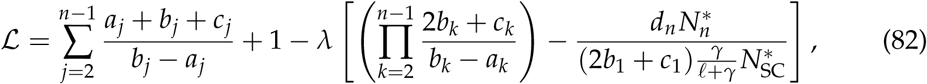

where we have used the expression for the steady-state population size from Eq. (4) in the main text to derive the constraint for the number of TDCs. As the parameters *b*_1_ (= *a*_1_), *c*_1_, *γ, ℓ*, and 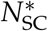 do not appear in the objective function (only in the constraint), they cannot be optimised here. Furthermore, the number of compartments *n* will be optimised through graphical methods due to its discrete nature.

Finding the stationary points of Eq. (82) is achieved by setting the derivatives *∂*ℒ/*∂p*_*i*_ (*p* ∈ {*a, b, c*}; 1 < *i* < *n*) and *∂*ℒ/*∂λ* equal to zero. These derivatives satisfy:

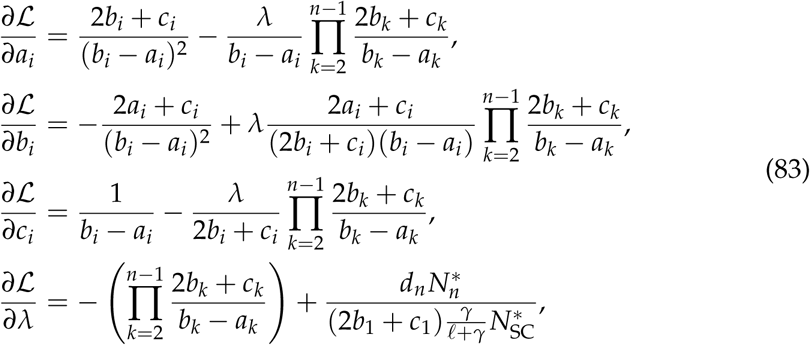

and hence we arrive at the four conditions:

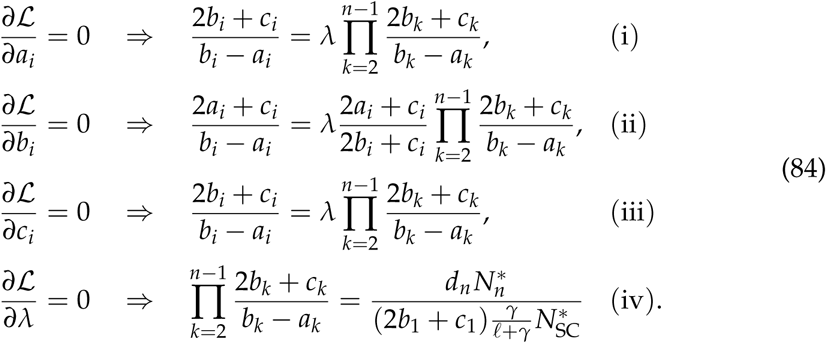

We have used the condition *b*_*i*_ − *a*_*i*_ > 0 (for 1 < *i* < *n*) above. We have also had to impose 2*b*_*i*_ + *c*_*i*_ > 0, which follows from the fact that we must have some differentiation flux from compartment *i* to *i* + 1 and therefore requires the condition *b*_*i*_ + *c*_*i*_ > 0 (as well as *b*_*i*_ ≥ 0 and *c*_*i*_ ≥ 0 by definition). The term 2*a*_*i*_ + *c*_*i*_ ≥ 0 can be zero iff *a*_*i*_ = 0 and *c*_*i*_ = 0, so we cannot divide by this term in general. The conditions (i) and (iii) above are equivalent, and condition (ii) can be shown to be consistent with (i) and (iii) by replacing the 2*b*_*i*_ + *c*_*i*_ with this value as defined by condition (i).

Ultimately, the conditions (i)-(iii) in Eq. (84) can be reduced to

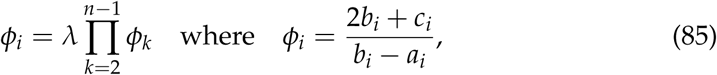

and hence the ratios *ϕ*_*i*_ are conserved across compartments 1 < *i* < *n*:

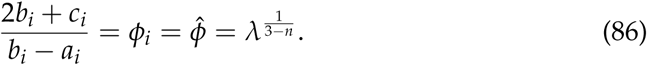

The value of *λ* is defined by condition (iv) in Eq. (84),

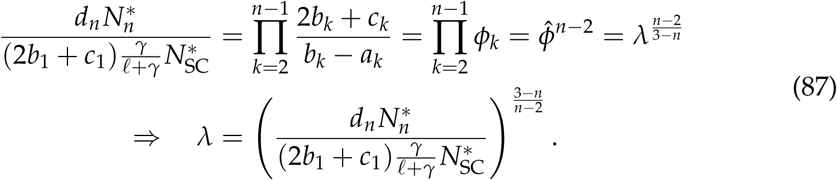

We can therefore define the optimal ratio 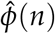 as

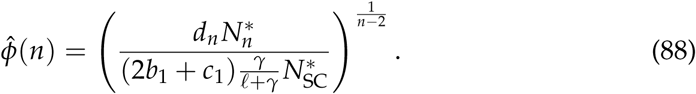

Inserting this value of 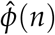 into the expression for the number of divisions per TDC after SC differentiation [Eq. (56)], we arrive at

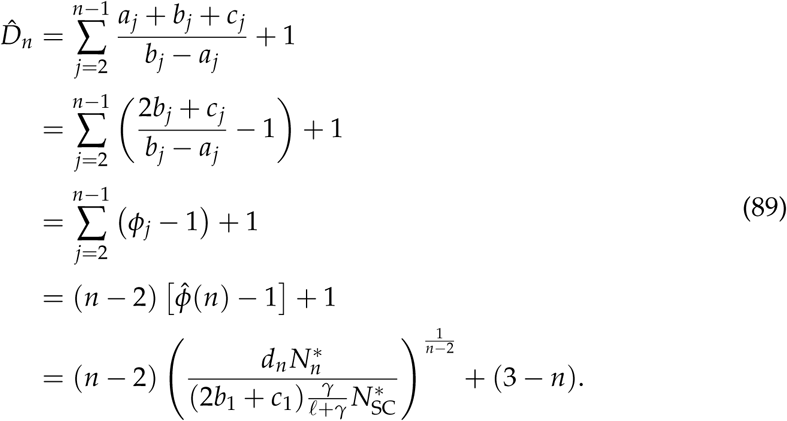

The divisional load is minimised when 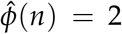. Because of the conditions *b*_*i*_ − *a*_*i*_ > 0, as well as *a*_*i*_ ≥ 0, *b*_*i*_ ≥ 0, and *c*_*i*_ ≥ 0, the value 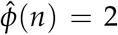 can only be achieved if *a*_*i*_ = *c*_*i*_ = 0, i.e. progenitor cells (1 < *i* < *n*) do not self-renew or divide asymmetrically, they only differentiate.

#### Minimizing cumulative divisions after lifetime *t*

As in the reduced model (Appendix III), we make the assumption that SC dynamics are much slower than non-SC dynamics, such that the number of divisions per TDC at time *t* is approximated by the number of SC divisions at time *t* plus the number of divisions that the TDC accumulates after leaving the SC compartment. I.e. the number of divisions per TDC at time *t* is

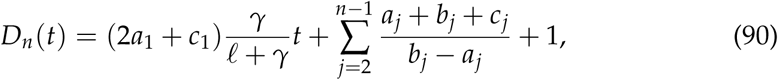

where we have already set *d*_*i*_ = 0 for 1 ≤ *i* < *n*. Here we minimise the number of divisions that TDCs will have accumulated after SCs have been dividing for a time period *t* = *τ*, including divisions accumulated in the stem cells up to this time. Note that *τ* is the timescale of optimisation, but the lifetime *t* may be longer or shorter than *τ*.

We now want to find the stationary points of the Lagrangian function after a lifetime *τ*:

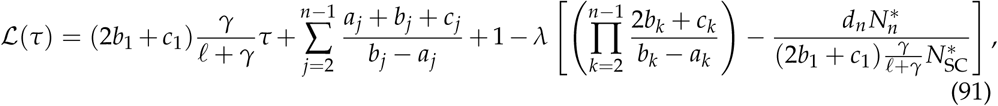

where we have used the expression for the steady-state population size from Eq. (4) to derive the constraint for the number of TDCs, as well as using *a*_1_ = *b*_1_ which is required for homeostasis.

Finding the stationary points of this Lagrangian function is achieved by setting the derivatives *∂*ℒ(*τ*)/*∂p*_*i*_ (*p* ∈ {*a, b, c*}), *∂*ℒ(*τ*)/*∂γ, ∂*ℒ(*τ*)/*∂ ℓ* and *∂*ℒ(*τ*)/*∂λ* equal to zero. These derivatives satisfy:

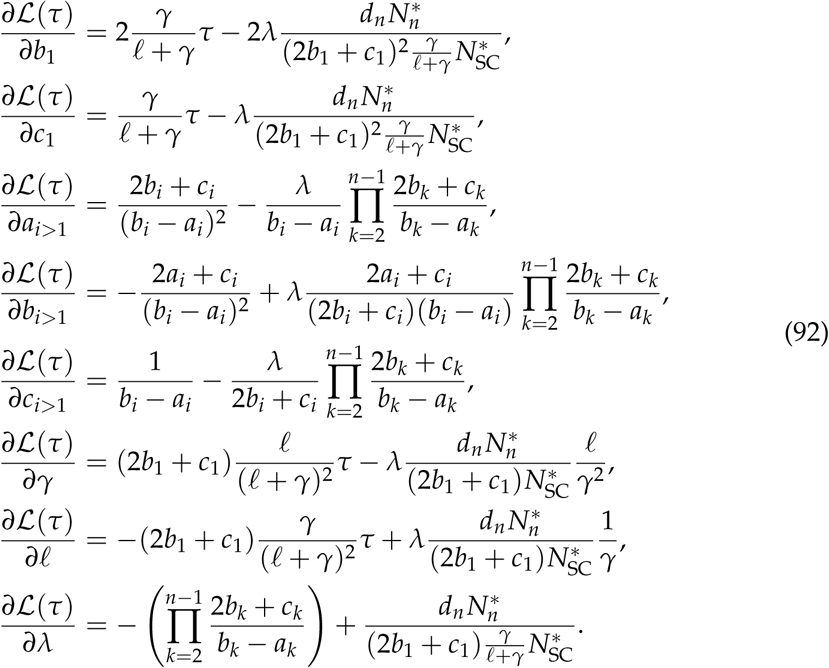

Equating each derivative to zero, we arrive at the following conditions:

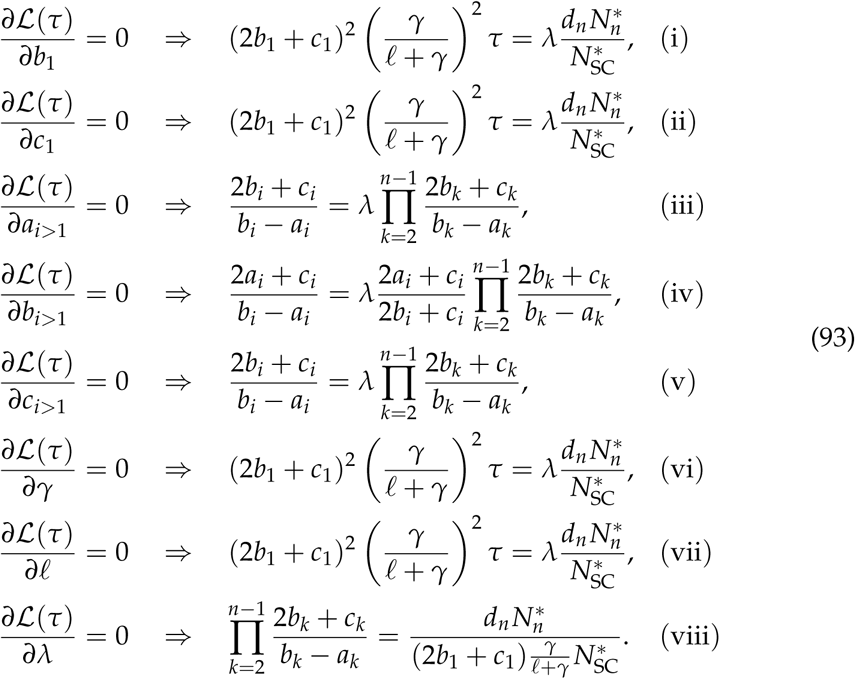

We have used the condition *b*_*i*_ − *a*_*i*_ > 0 above. We have also had to impose 2*b*_*i*_ + *c*_*i*_ > 0, which follows from the fact that we must have some differentiation flux from compartment *i* to *i* + 1 and therefore requires the condition *b*_*i*_ + *c*_*i*_ > 0 (as well as *b*_*i*_ ≥ 0 and *c*_*i*_ ≥ 0 by definition). The term 2*a*_*i*_ + *c*_*i*_ ≥ 0 can be zero iff *a*_*i*_ = 0 and *c*_*i*_ = 0, so we cannot divide by this term in general. The conditions (iii) and (v) above are equivalent, and condition (iv) can be shown to be consistent with (iii) and (v) by replacing the 2*b*_*i*_ + *c*_*i*_ with this value as defined by condition (iii). Meanwhile, conditions (i), (ii), (vi), and (vii) are equivalent.

Ultimately, the conditions (iii)-(v) in Eq. (93) can be reduced to

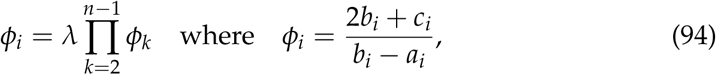

and hence the ratios *ϕ*_*i*_ are conserved across compartments 1 < *i* < *n*:

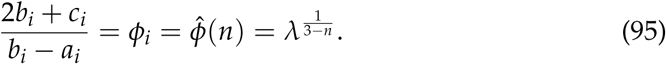

Conditions (i), (ii), (vi), and (vii) in Eq. (93) can be reduced to

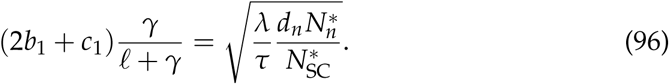

From condition (viii), i.e. the steady state population size, the value of *λ* is then defined as

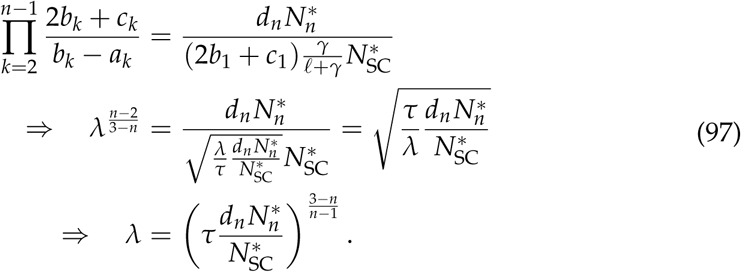

Therefore, the optimised parameter combinations satisfy

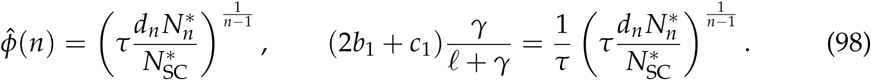

The cumulative number of divisions after time 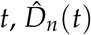, then satisfies

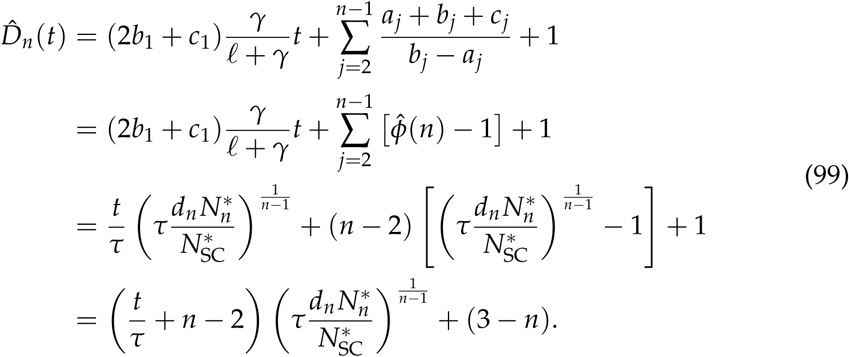

